# HIF-1α induces glycolytic reprogramming in tissue-resident alveolar macrophages to promote survival during acute lung injury

**DOI:** 10.1101/2022.02.28.482301

**Authors:** Parker S. Woods, Lucas M. Kimmig, Kaitlyn A. Sun, Angelo Y. Meliton, Obada R. Shamaa, Yufeng Tian, Rengül Cetin-Atalay, Willard W. Sharp, Robert B. Hamanaka, Gökhan M. Mutlu

**Affiliations:** Department of Medicine, Section of Pulmonary and Critical Care Medicine, The University of Chicago, Chicago, IL 60637; Department of Medicine, Section of Emergency Medicine, The University of Chicago, Chicago, IL 60637

## Abstract

Cellular metabolism is a critical regulator of macrophage effector function. Tissue-resident alveolar macrophages (TR-AMs) inhabit a unique niche marked by high oxygen and low glucose. We have recently shown that in contrast to bone marrow-derived macrophages (BMDMs), TR-AMs cannot utilize glycolysis and predominantly rely on mitochondrial function for their effector response. It is not known how changes in local oxygen concentration that occur during conditions such as acute respiratory distress syndrome (ARDS) might affect TR-AM metabolism and function; however, ARDS is associated with progressive loss of TR-AMs, which correlates with the severity of disease and mortality. Here, we demonstrate that hypoxia robustly stabilizes HIF-1α in TR-AMs to promote a glycolytic phenotype. Hypoxia altered TR-AM metabolite signatures, cytokine production, and decreased their sensitivity to the inhibition of mitochondrial function. By contrast, hypoxia had minimal effects on BMDM metabolism. The effects of hypoxia on TR-AMs were mimicked by FG-4592, a HIF-1α stabilizer. Treatment with FG-4592 decreased TR-AM death and attenuated acute lung injury in mice. These findings reveal the importance of microenvironment in determining macrophage metabolic phenotype, and highlight the therapeutic potential in targeting cellular metabolism to improve outcomes in diseases characterized by acute inflammation.

## INTRODUCTION

Glycolytic metabolism has been ascribed a central role in macrophage inflammatory processes (Tannahill, Curtis et al. 2013, Freemerman, Johnson et al. 2014, Palsson-McDermott, Curtis et al. 2015, Xie, Yu et al. 2016, Ip, Hoshi et al. 2017). Much of our current understanding of this phenomena has been elucidated in bone marrow-derived macrophages (BMDMs) and macrophage cell lines (i.e., THP-1 and RAW 264.7), which model macrophages of monocytic lineage. Considerably less is known about how other factors like local microenvironment and developmental origin may influence macrophage metabolic function. Tissue-resident alveolar macrophages (TR-AMs) reside within the lumen of the lung alveolus where they are critical in maintaining lung homeostasis within the healthy airway and are the first responders to airborne pathogens and pollutants (Hussell and Bell 2014). The alveolus maintains the highest oxygen concentration of any tissue compartment within the human body (Carreau, El Hafny-Rahbi et al. 2011). Moreover, under steady-state conditions, glucose concentrations within airway lumen are less than one-tenth of blood glucose concentrations (Baker, Clark et al. 2007). These environmental conditions alone suggest a requirement for oxidative metabolism for cells that reside within the alveoli. Several studies have demonstrated that the unique characteristics of the alveolar microenvironment heavily influence macrophage function and immunometabolism (Lavin, Winter et al. 2014, Svedberg, Brown et al. 2019, McQuattie-Pimentel, Ren et al. 2021). Our group has recently demonstrated that TR-AMs rely predominantly on oxidative phosphorylation under steady-state conditions and that glycolysis is dispensable for proinflammatory effector function in these cells (Woods, Kimmig et al. 2020). Together, these findings highlight the lung microenvironment’s central role in dictating TR-AM responses.

Conditions associated with severe airway inflammation (i.e., acute respiratory distress syndrome (ARDS)) increase alveolar epithelial/endothelial barrier permeability (Ware and Matthay 2000). This results in flooding of alveoli with fluid and recruitment of non-resident immune cells leading to severe local hypoxia as well as an increase in alveolar glucose levels (Fröhlich, Boylan et al. 2013, Campbell, Bruyninckx et al. 2014, Baker and Baines 2018). These abrupt changes in the alveolar microenvironment under a diseased state such as ARDS likely necessitate metabolic adaptation by TR-AMs in order to ensure optimal cellular fitness. Acute lung injury/ARDS is associated with a decline in the number of TR-AMs and the degree of TR-AM loss correlates with clinical outcomes (i.e., mortality) (Fan and Fan 2018). Moreover, experimental depletion of TR-AMs results in an increase in the severity of acute lung injury and mortality (Beck-Schimmer, Schwendener et al. 2005, Kim, Lee et al. 2008, Jaworska, Coulombe et al. 2014, Machado-Aranda, V Suresh et al. 2014, Nelson, Zhou et al. 2014, Schneider, Nobs et al. 2014, Cardani, Boulton et al. 2017). Whether or not changes in the alveolar microenvironment play a role in TR-AM cell death during acute lung injury/ARDS has yet to be explored.

Hypoxia-inducible factor 1-alpha (HIF-1α) is the most extensively characterized transcription factor responsible for cellular adaptation to low oxygen levels (Semenza 2012). In brief, under normoxia, oxygen-dependent proline hydroxylases prevent HIF-1α activation by marking it for proteasomal degradation. Conversely, under hypoxia, decreased hydroxylase activity promotes HIF-1α protein stabilization and translocation to the nucleus allowing for transcriptional responses to low oxygen levels, such as enhanced expression of genes related to glycolysis and angiogenesis. HIF-1α has been well characterized in macrophages of monocytic origin, and identified to play a key role in macrophage infiltration and proinflammatory responses (Cramer, Yamanishi et al. 2003, Peyssonnaux, Datta et al. 2005, Tannahill, Curtis et al. 2013, Matak, Heinis et al. 2015, Palsson-McDermott, Curtis et al. 2015). However, little is known about the role that HIF-1α plays in TR-AM effector function and metabolism. In a study focusing on TR-AM development, Izquierdo *et al* demonstrated that the expression of HIF-1α and its target genes is turned off following birth and this process is required for TR-AM maturation and normal effector function (Izquierdo, Brandi et al. 2018). It is unknown; however, whether HIF-1α plays a role in TR-AM effector function after maturation or how local hypoxia and changes in the expression of HIF-1α might regulate the adaptation of mature TR-AMs to hypoxia or affect their effector function during acute lung injury.

To answer these questions, we used a variety of metabolic and immunological approaches. We observed that HIF-1α was undetectable in primary TR-AMs cultured under normoxia, but was robustly stabilized under hypoxia in a dose-dependent fashion. Upon hypoxic HIF-1α stabilization, TR-AMs acquired a glycolytic phenotype, which was not observed under normoxic conditions. In contrast, BMDMs exhibited no alterations in HIF-1α stabilization or glycolytic output in response to hypoxia. Analysis of LPS-induced TR-AM metabolite signatures revealed significant increases in glycolytic intermediates under hypoxia. Hypoxia also altered TR-AM cytokine profile in response to LPS. In contrast to BMDMs, TR-AMs depend on mitochondrial function for inflammatory responses under normoxia. Hypoxia rescued the ETC inhibitor-induced impairment in cytokine production in TR-AMs.

Using influenza infection in mice to model acute lung injury, we found that TR-AM cell number decreased over the course of acute lung injury. Surviving TR-AMs exhibited a glycolytic gene signature supporting hypoxic adaptation. To determine how the ability to adapt to hypoxia affects TR-AM survival and function, we treated influenza-infected mice intratracheally with FG-4592, a HIF-1α stabilizer, which mimics hypoxic adaptation. Compared to control mice, FG-4592 prevented the loss of TR-AMs, reduced lung injury, and increased survival. Collectively, our data suggest that HIF-1α plays a critical role in TR-AM metabolic adaptation to altered environmental conditions during acute lung injury. Promoting hypoxic adaptation and glycolytic metabolism in TR-AMs enables them to adapt to and survive the changes in the microenvironment during lung injury and consequently reduces lung inflammation and may offer a viable therapeutic strategy in treating ARDS arising from influenza or other severe viral infections, including COVID19.

## RESULTS

### Tissue-resident alveolar macrophages exhibit HIF-1α stabilization and develop a glycolytic phenotype in response to hypoxia

We have recently shown that TR-AMs maintain a very low glycolytic rate which is not augmented by activation of inflammatory responses (Woods, Kimmig et al. 2020). As TR-AMs inhabit an environment with high oxygen levels, we hypothesized that TR-AMs may not be able to induce glycolytic reprogramming in response to either inflammatory stimuli or to physiologic hypoxia. Glycolysis stress tests were performed following overnight (16 hours) exposure to decreasing levels of ambient oxygen. Unlike the inability of TR-AMs to induce glycolysis after inflammatory stimulus under normoxia, they exhibited a progressive increase in the extracellular acidification rate (ECAR) in response to escalating degrees of ambient hypoxia (Figure 1A). Both basal rate of glycolysis and glycolytic reserve increased substantially when oxygen levels were lowered to 3.0% and 1.5% (Figure 1B). HIF-1α levels were nearly undetectable under normoxic conditions; however, with increasing degrees of hypoxia, HIF-1α stabilization occurred in a dose-dependent fashion and was detectable in the nucleus (Figure 1C). Intracellular lactate levels increased slightly in TR-AMs after exposure to hypoxia compared with normoxic control cells while the lactate levels in the media increased by a greater degree (Figure 1D). Pretreating TR-AMs prior to hypoxia with echinomycin, an inhibitor of HIF-1α DNA binding activity (Kong, Park et al. 2005), disrupted hypoxia-induced increases in glycolytic output in a dose-dependent fashion (Figure 1E). Echinomycin also reduced hypoxia-induced increases in glycolytic protein expression (LDHA and HK2), suggesting that HIF-1α is required for glycolytic adaption to hypoxia in TR-AMs (Figure 1F).

**Figure 1.**
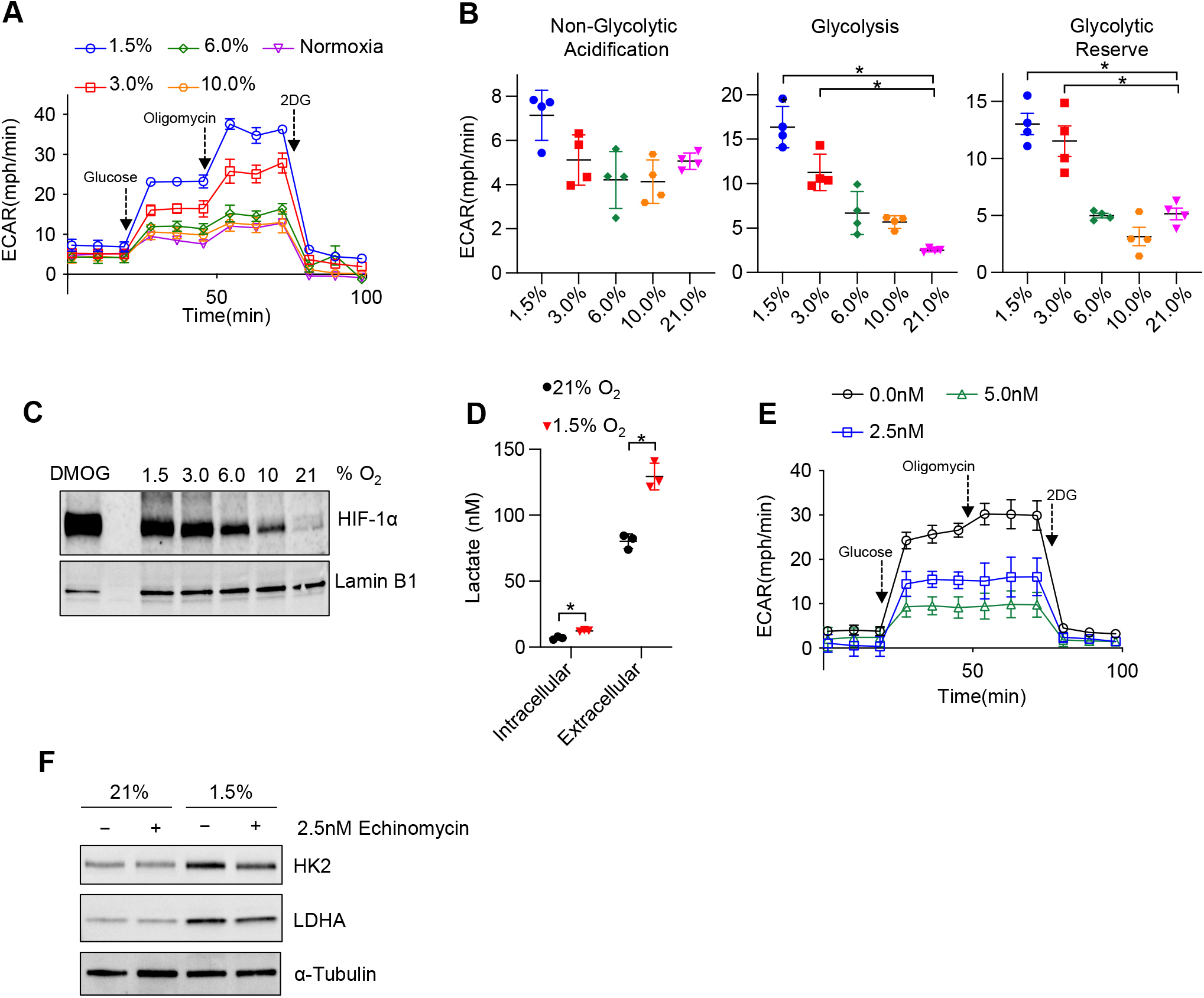
Tissue-resident alveolar macrophages exhibit HIF-1α stabilization and develop a glycolytic phenotype in response to hypoxia. TR-AMs were incubated overnight (16h) at varying O_2_ concentrations. **(A)** Using Seahorse XF24 analyzer, glycolysis was measured as extracellular acidification rate (ECAR). TR-AMs were sequentially treated with glucose, oligomycin (ATP synthase inhibitor) and 2-deoxyglucose (2-DG) (inhibitor of hexokinase 2, or glycolysis). **(B)** Interleaved scatter plots quantifying glycolytic parameters. Data represent at least 3 independent experiments (n=4 separate wells per group). Glycolytic parameters were compared against 21% O_2_ and significance was determined by one-way ANOVA with Bonferroni’s post test. **(C)** Western blot analysis of nuclear extract to assess HIF-1α expression. DMOG served as a positive control **(D)** Extracellular and intracellular lactate levels in TR-AMs incubated overnight (16h) under normoxia or 1.5% O_2_. Significance was determined by two-way ANOVA with Bonferroni’s post test. **(E)** Glycolysis stress test of TR-AMs under 1.5% O_2_ in combination with echinomycin (16h). **(F)** Western blot analysis of whole cell TR-AM extract after 16h treatment. All error bars denote mean ± SD. *, p < 0.05

Both short-term (2 hours) and prolonged (16 hours) exposure to hypoxia (1.5% O_2_) led to significant increases in nuclear HIF-1α protein levels in TR-AMs (Figure S1A). Glycolysis stress tests demonstrated that short-term hypoxia treatment failed to induce significant alterations in glycolysis or glycolytic capacity in TR-AMs compared to prolonged hypoxia treatment suggesting that transcription and translation of glycolytic genes that are targets of HIF-1α are required (Figure S1B,C). Taken together, these data indicate that TR-AM HIF-1α stabilization in response to hypoxia is dose-dependent, and that prolonged hypoxia, but not short-term hypoxia exposure, leads to a functional glycolytic phenotype in TR-AMs.

### BMDMs have limited metabolic adaptation to hypoxia

Several studies have examined the effects of hypoxia on BMDM metabolism; however, they focused heavily on transcriptional changes in glycolytic gene expression as opposed to functional changes in glycolysis (Bosco, Puppo et al. 2006, Roiniotis, Dinh et al. 2009, Delprat, Tellier et al. 2020). We found that, unlike TR-AMs, BMDMs exposed to hypoxia (16 hours) exhibit minimal changes in glycolytic rate or glycolytic capacity (Figure 2A–B). Interestingly, we found that BMDMs have high basal levels of nuclear HIF-1α protein under normoxic conditions and that HIF-1α expression in BMDMs did not significantly change in response to hypoxia as low as 1.5% O_2_ (Figure 2C). Exposure to hypoxia resulted in only a small increase in extracellular lactate levels (Figure 2D), which is in agreement with the glycolysis stress test data. Echinomycin had a minimal effect on the glycolytic output of hypoxic BMDMs (Figure 2E). Likewise, neither hypoxia nor hypoxia in combination with echinomycin altered glycolytic protein expression (LDHA and HK2) in BMDMs (Figure 2F). Duration of hypoxia exposure (2h vs 16h) had no significant effect on HIF-1α stabilization or glycolysis in BMDMs (Figure S2A–C). Collectively, these data demonstrate that hypoxia has minimal effect on glycolytic function and HIF-1α stabilization in BMDMs.

**Figure 2.**
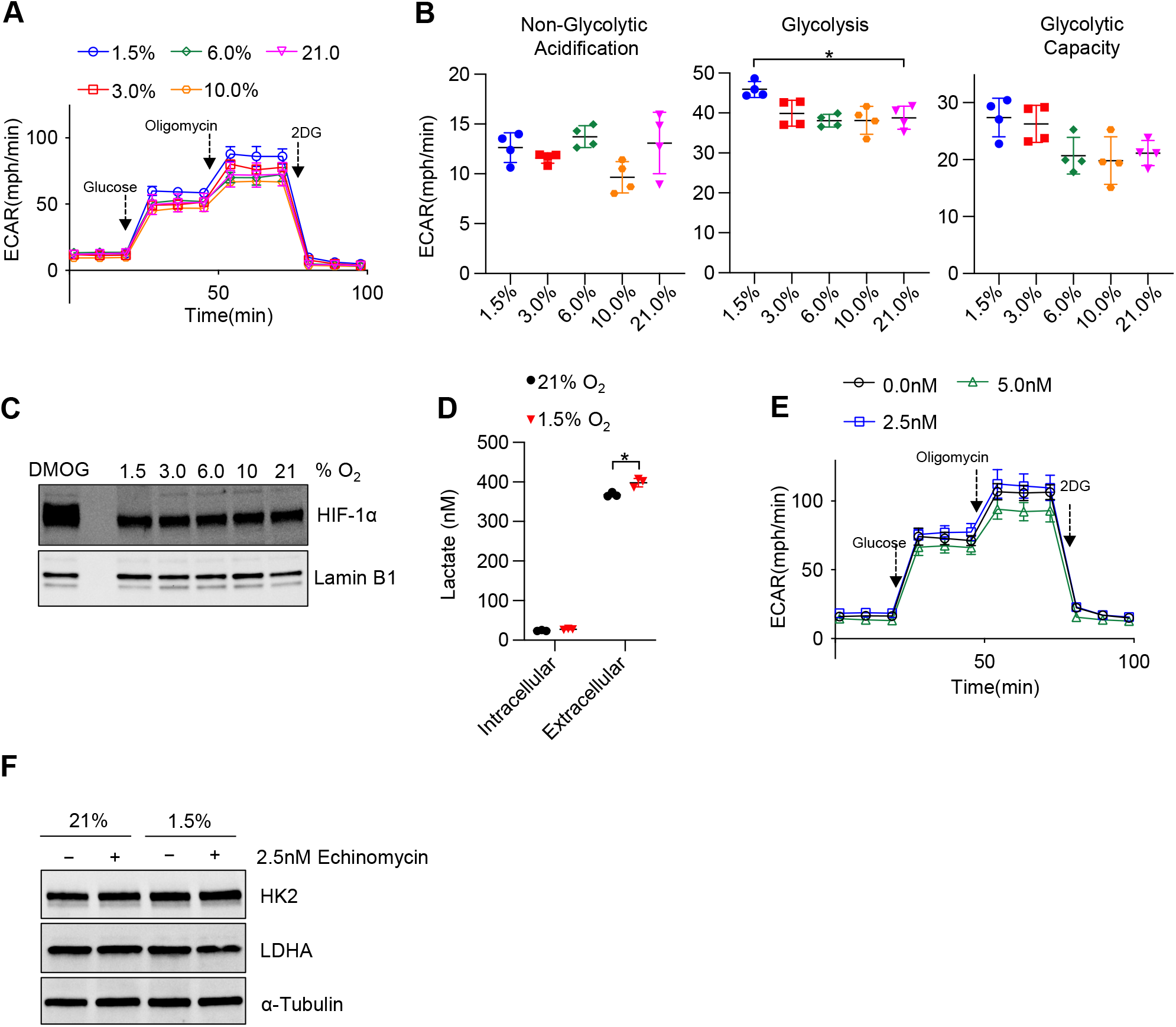
BMDMs have limited metabolic adaptation to hypoxia. BMDMs were incubated overnight (16h) at varying O_2_ concentrations. **(A)** Using Seahorse XF24 technology, glycolysis was measured as extracellular acidification rate (ECAR). **(B)** Interleaved scatter plots quantifying glycolytic parameters. Data represent at least 3 independent experiments (n=4 separate wells per group). Glycolytic parameters were compared against 21% O_2_ and significance was determined by one-way ANOVA with Bonferroni’s post test. **(C)** Western blot analysis of nuclear extracts to assess HIF-1α expression. **(D)** Extracellular and intracellular lactate levels in BMDMs incubated overnight (16h) under 21% or 1.5% O_2_. Significance was determined by two-way ANOVA. with Bonferroni’s post test. **(E)** Glycolysis stress test of BMDMs under 1.5% O_2_ in combination with echinomycin (16h). **(F)** Western blot analysis of whole cell extract from BMDMs treated with normoxia (21% O_2_) or hypoxia (1.5% O_2_) for 16h. All error bars denote mean ± SD. *, p < 0.05

### The hypoxia-induced transcriptomic response differs substantially between TR-AMs and BMDMs

To better understand the observed differences in hypoxia-induced glycolytic metabolism between TR-AMs and BMDMs, we performed RNA-sequencing to assess global alterations in gene expression. We found 741 DEGs (512 upregulated and 229 downregulated) in TR-AMs in response to 1.5% O_2_ compared to only 260 DEGs (214 upregulated and 46 downregulated) in BMDMs (Figure 3A). Reactome pathway analysis revealed that hypoxia altered a large number of TR-AM genes in multiple pathways ranging from cellular metabolism, hemostasis, and immune cell function, while the majority of BMDM genes affected by hypoxia were related to carbohydrate metabolism (Figure 3B). Hypoxia led to the most significant increases in glycolytic and HIF-1α regulatory gene expression in both TR-AMs and BMDMs. These same genes were significantly lower in TR-AMs compared to BMDMs under normoxic conditions (Figure 3C). This is in direct agreement with our previous findings (Woods, Kimmig et al. 2020). A side-by-side comparison demonstrated that the level of HIF-1α expression in hypoxic TR-AMs is similar to that of BMDMs under normoxia and hypoxia (Figure 3D). Hypoxia-induced HIF-1α expression in TR-AMs correlated with increase in glycolytic (HK2 and LDHA) and prolyl hydroxylase (EGLN1 and EGLN3) protein expression in TR-AMs (Figure 3E). This was not the case in BMDMs in which hypoxia exposure did not alter protein expression of glycolytic genes. These results demonstrate that hypoxia induces transcriptomic alterations in TR-AMs that lead to changes in protein expression and metabolic function. In contrast, BMDMs exhibit high levels of HIF-1α and HIF-1α target proteins at baseline, and do not further increase the expression of these proteins in response to hypoxia, despite a hypoxia-adaptive mRNA transcription (Figure 3D and 3E).

**Figure 3.**
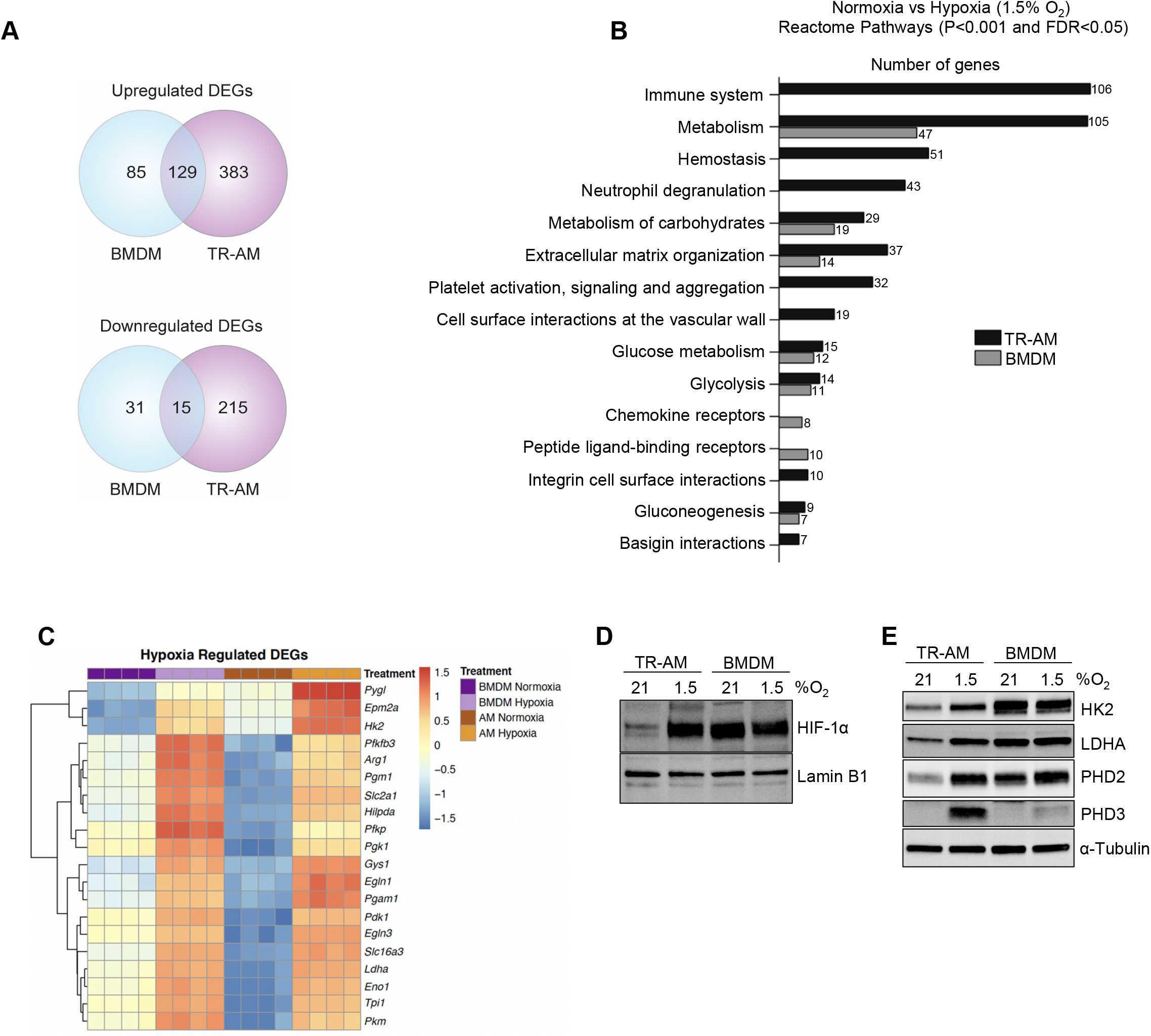
The hypoxia-induced transcriptomic response differs substantially between TR-AMs and BMDMs. TR-AMs and BMDMs were incubated overnight (16h) under normoxia (21.0% O_2_) or hypoxia (1.5% O_2_). **(A)** Venn diagrams show differentially expressed genes (DEGs) altered by hypoxia in TR-AMs (741 total DEGs), and BMDMs (260 total DEGs). DEGs were identified using DESeq2 at FC>2 and FDR adjusted p-value of p<0.05. **(B)** Reactome pathway enrichment comparing number of genes in a given pathway altered by hypoxia in TR-AMs and BMDMs. **(C)** Heatmap representing the top 20 significant metabolic genes altered by hypoxia in both TR-AMs and BMDMs. **(D)** Western blot analysis of nuclear extracts to assess HIF-1α protein expression. (**E)** Western blot analysis of whole cell extracts to assess glycolytic enzyme (HK2, LDH) and prolyl hydroxylase (PHD2, PHD3) protein expression.

### Hypoxia modulates TR-AM cytokine production and metabolic response to LPS

Hypoxia and HIF-1α are thought to be central to the inflammatory response of macrophages (Cramer, Yamanishi et al. 2003, Tannahill, Curtis et al. 2013, Palsson-McDermott, Curtis et al. 2015). To determine the effect of HIF-1α stabilization on TR-AM’s effector response, we measured the production of proinflammatory cytokines in response to LPS under hypoxia. TR-AMs were exposed overnight to hypoxia (1.5% O_2_) or normoxia and then subsequently treated with LPS while maintaining original O_2_ conditions. Hypoxia alone did not stimulate cytokine production without LPS treatment. Hypoxic TR-AMs secreted significantly higher levels of TNF-α, KC, and IL-1β in response to LPS compared to normoxic controls. In contrast, IL-6 and CCL2 secretion was decreased in hypoxic TR-AMs (Figure 4A). The cytokine gene expression pattern in hypoxic TR-AMs treated with LPS mirrored the secreted cytokine profile (Figure 4B). Moreover, enhanced proIL-1β protein production was observed in hypoxic TR-AMs treated with LPS (Figure 4C). While BMDMs experienced limited metabolic alterations in response to hypoxia, treatment with LPS revealed that hypoxia induced similar alterations in their cytokine profile. Hypoxic BMDMs had increased TNF-α, KC, and IL-1β, and decreased IL-6 secretion (Figure S3A). The only discordance in the hypoxic cytokine response profile between TR-AMs and BMDMs was CCL2, which remained unchanged in hypoxic BMDMs in response to LPS compared to normoxic controls (Figure S3A).

**Figure 4.**
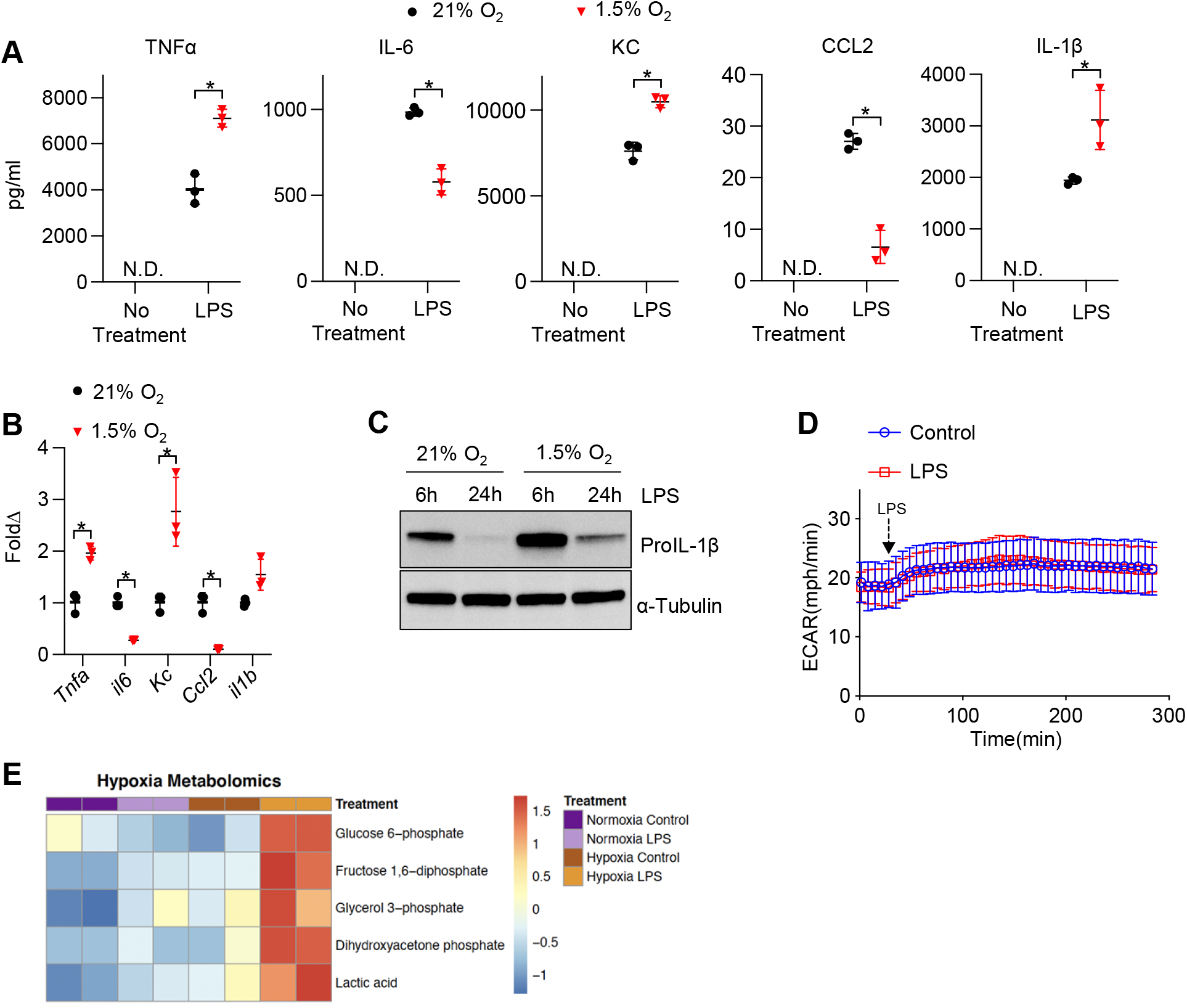
Hypoxia modulates TR-AM cytokine production and metabolic response to LPS. TR-AMs were incubated overnight (16h) under 21% or 1.5% O_2_ then stimulated with 20ng/ml LPS for 6 while maintaining pretreatment conditions. For IL-1β measurements, 5mM ATP was added to TR-AMs for 30 minutes following 6h LPS treatment to activate caspase 1, ensuring IL-1β release. **(A)** We measured cytokine (TNFα, IL-6, KC, CCL2 and IL-1β) levels in media using ELISA. Data represent at least 3 independent experiments; n=3 per group. Significance was determined by two-tailed T Test **(B)** qPCR was used to measure mRNA expression (*Tnfa, il6, Kc, Ccl2, and il1b*). Gene expression was normalized to corresponding gene ct values in 21% group and represented as fold change using the ΔΔct method. Data represent at least 3 independent experiments; n=3 per group. Significance was determined by two-tailed T Test **(C)** Western blot analysis of whole cell extracts at 6 and 24h post LPS treatment **(D)** ECAR was measured in following acute LPS injection (final concentration: 20 ng/ml) in TR-AMs conditioned in 1.5% O_2_ **(E)** CE-MS metabolite heatmap for glycolytic intermediates. All error bars denote mean ± SD. *, p < 0.05.

We and others have shown that BMDMs exhibit an immediate enhancement in glycolytic output in response to LPS (Figure S4). It is thought that this increase in glycolysis following LPS supports the proinflammatory response. We have shown that LPS-induced inflammation in TR-AMs is independent of glycolysis including the rise in glycolysis following LPS injection (Woods, Kimmig et al. 2020). Given that hypoxia elevated HIF-1α levels and glycolytic rates in TR-AMs, we sought to determine if hypoxia could alter TR-AM glycolytic responsiveness to LPS. We found that despite the fact that hypoxia increased the glycolytic rate of TR-AMs at baseline, TR-AM glycolysis remained unresponsive to LPS injection (Figure 4D). Using capillary electrophoresis-mass spectrometry to measure glycolytic metabolite levels, we found that consistent with increased glycolytic output after hypoxia, levels of glycolytic intermediate metabolites (glucose-6 phosphate, fructose 1,6 diphosphate, glycerol 3-phosphate, dihydroxyacetone phosphate) and lactate were increased in response to hypoxia alone. Interestingly, hypoxic TR-AMs exhibited further increases in glycolytic intermediates in the presence of LPS (6h) compared to normoxic cells, suggesting that while LPS increases cellular levels of glycolytic metabolites in hypoxic TR-AMs, this does not manifest as acute lactate secretion (Figure 4E). These data demonstrate that hypoxia leads to significant alterations in TR-AM cytokine production and increased glycolytic metabolites in response to prolonged LPS treatment. However, unlike the prototypical BMDM response, hypoxic TR-AMs do not immediately increase their extracellular acidification in response to LPS.

We have previously shown that unlike BMDMs, TR-AMs effector function is acutely sensitive to mitochondrial inhibition (Woods, Kimmig et al. 2020). Since TR-AM capacity for glycolysis expands with decreasing levels of O_2_, we next sought to assess mitochondrial function under hypoxia and performed a mitochondrial stress test on TR-AMs that had been exposed to varying oxygen concentrations. Interestingly, mild-to-moderate degrees of ambient hypoxia did not appear to significantly alter overall mitochondrial function in these cells. Only 1.5% O_2_ caused significant reductions in oxygen consumption rate (OCR) across all mitochondrial parameters (Figure 5A, B). ECAR tracings during the mitochondrial stress test demonstrated that, other than severe hypoxia (1.5% O_2_), the majority of acid produced under mild-moderate hypoxia is derived from CO_2_, as the application of rotenone and antimycin A led to a significant reduction in ECAR (Figure 5C). When exposed to 1.5% O_2_, TR-AM energy is derived mostly from glycolysis with little TCA activity and no significant contribution of CO_2_ to extracellular acidification. Overall BMDM mitochondrial function was impaired by 1.5% O_2_, but the effect was greatly diminished compared to TR-AMs (Figure S5A, B). BMDM ECAR tracing during mitochondrial stress test demonstrated that most acid production remained unchanged in response to rotenone and antimycin A regardless of O_2_ concentration (Figure S5C). This suggests that BMDM acidification is glycolytically-derived under both normoxia and hypoxia.

**Figure 5.**
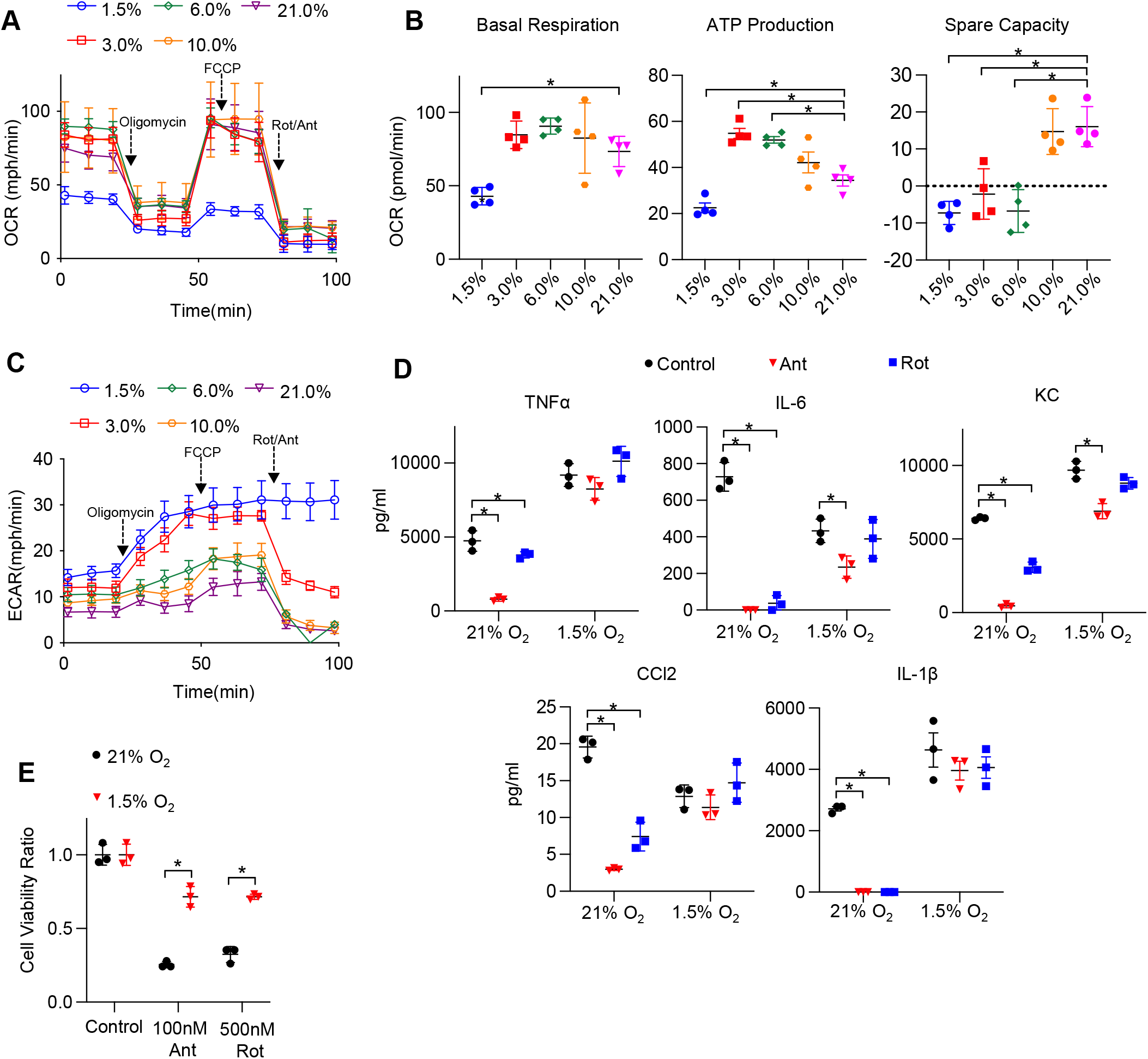
Hypoxia rescues ETC inhibitor-induced cell death and impaired cytokine production in TR-AMs. **(A)** Mitochondrial stress test to measure oxygen consumption rate (OCR) using Seahorse XF24 in TR-AMs, which were treated sequentially with oligomycin (ATP synthase inhibitor), FCCP (uncoupler) and Rotenone (Rot)/Antimycin A (Ant) (complex I and III inhibitors, respectively). **(B)** Interleaved scatter plots quantifying mitochondrial respiration parameters. Data represents at least 3 experiments (n=4 separate wells per group). Mitochondrial parameters were were compared against 21% O_2_ and significance was determined by one-way ANOVA with Bonferroni’s post test **(C)** ECAR measurement during mitochondrial stress test to visualize TR-AMs ability to upregulate glycolysis in response to mitochondrial inhibition **(D)** TR-AMs were incubated overnight (16h) under 21% or 1.5% O_2_ then stimulated with 20ng/ml LPS in the presence of absence of mitochondrial inhibitors (20nM Ant or Rot) for 6 hours while maintaining pretreatment conditions. ELISA was used to measure secreted cytokine (TNFα, IL-6, KC, CCL2, and IL-1β) levels in media. ATP added to cells prior to collection for IL-1β assessment. Data represent at least 3 independent experiments; n=3 per group. Significance was determined by one-way ANOVA with Bonferroni’s post test. **(E)** TR-AMs were cultured under 21% or 1.5% O_2_ for 6h then treated with mitochondrial inhibitors (100nM Ant or 500nM Rot) overnight and an Sulforhodamine B assay was performed to measure cytotoxicity. Graphs represent cell viability compared to control, 21% O_2_ group. Data represent at least 3 independent experiments (n=3 per group). Significance was determined by two-way ANOVA with Bonferroni’s post test. All error bars denote mean ± SD. *, p < 0.05.

TR-AM cytokine production in response to LPS was highly susceptible to inhibition by low doses of ETC inhibitors, rotenone and antimycin A, under normoxic conditions. This effect was greatly attenuated after exposure to hypoxia (Figure 5D). Additionally, high doses of ETC inhibitors induce cytotoxicity in normoxic TR-AMs, but hypoxic preconditioning significantly enhanced TR-AM cell viability (Figure 5E). In contrast, BMDM cytokine production was only marginally affected by ETC inhibition with the exception of observed decrease in IL-1β. Unlike TR-AMs, hypoxia did not significantly alter BMDM cytokine production in the presence of ETC inhibitors (Figure S5D). Similarly, ETC inhibition did not to induce cytotoxicity in BMDMs under normoxia or hypoxia (Figure S5E).

### TR-AM survival correlates with a shift to glycolytic metabolism during influenza-induced acute lung injury

LPS is a well-known and potent macrophage activator that in isolation can be used to investigate essential immune functions, such as cytokine production, signal transduction, and immunometabolism. It induces a broad range of inflammatory effects in macrophages making it convenient tool to study overall immune fitness *in vitro.* However, *in vivo* studies have shown that LPS instillation into the murine airway leads to an immune response predominated by infiltrating neutrophils making LPS-induced ALI an unsuitable model to study macrophages (Chignard and Balloy 2000). Compared to LPS, influenza infection is a more clinically relevant model of ARDS, and various macrophages populations play a larger role in both exacerbating and limiting lung injury in this model (Short, Kroeze et al. 2014). Several groups have demonstrated that TR-AMs undergo cell death in response to influenza infection and that depletion of TR-AMs is associated with worse outcomes in models of influenza-induced ALI (Kim, Lee et al. 2008, Jaworska, Coulombe et al. 2014, Nelson, Zhou et al. 2014, Schneider, Nobs et al. 2014, Cardani, Boulton et al. 2017). To confirm this phenomenon, we utilized PKH26 Red Fluorescent Linker dye to specifically label, track, and collect TR-AMs over the time course of infection as we and others have previously described (Maus, Herold et al. 2001, Maus, Grote et al. 2002, Woods, Kimmig et al. 2020). In agreement with Zhu et al. (Zhu, Wu et al. 2021), we found that there was a significant decrease in TR-AMs (PKH26+) at 3 (D3) and 6 (D6) days post infection along with a subsequent increase in infiltrating, monocyte-derived alveolar macrophages (Mo-AMs) (Figure 6A). From these experiments, we performed RNAseq on sorted TR-AMs (PKH26+) and Mo-AMs (PKH26−) at D0, D3, and D6 (note: Mo-AMs are not present in an uninfected (D0) mouse)) to identify changes in the metabolic gene signature of these two macrophage populations during influenza infection. From D0 to D6, RNAseq data revealed that TR-AMs experienced decreased expression in genes related to oxidative phosphorylation with simultaneous increased expression of genes related to glycolytic metabolism. Moreover, the metabolic gene signature of D6 TR-AMs was most similar to that of Mo-AMs at D3 and D6 (Figure 6B, C). Taken together, these data suggest that influenza-induced ALI leads to a decrease in TR-AM number, and that the surviving TR-AMs’ gene signature shifts away from genes related to mitochondrial metabolism in favor of glycolysis.

**Figure 6.**
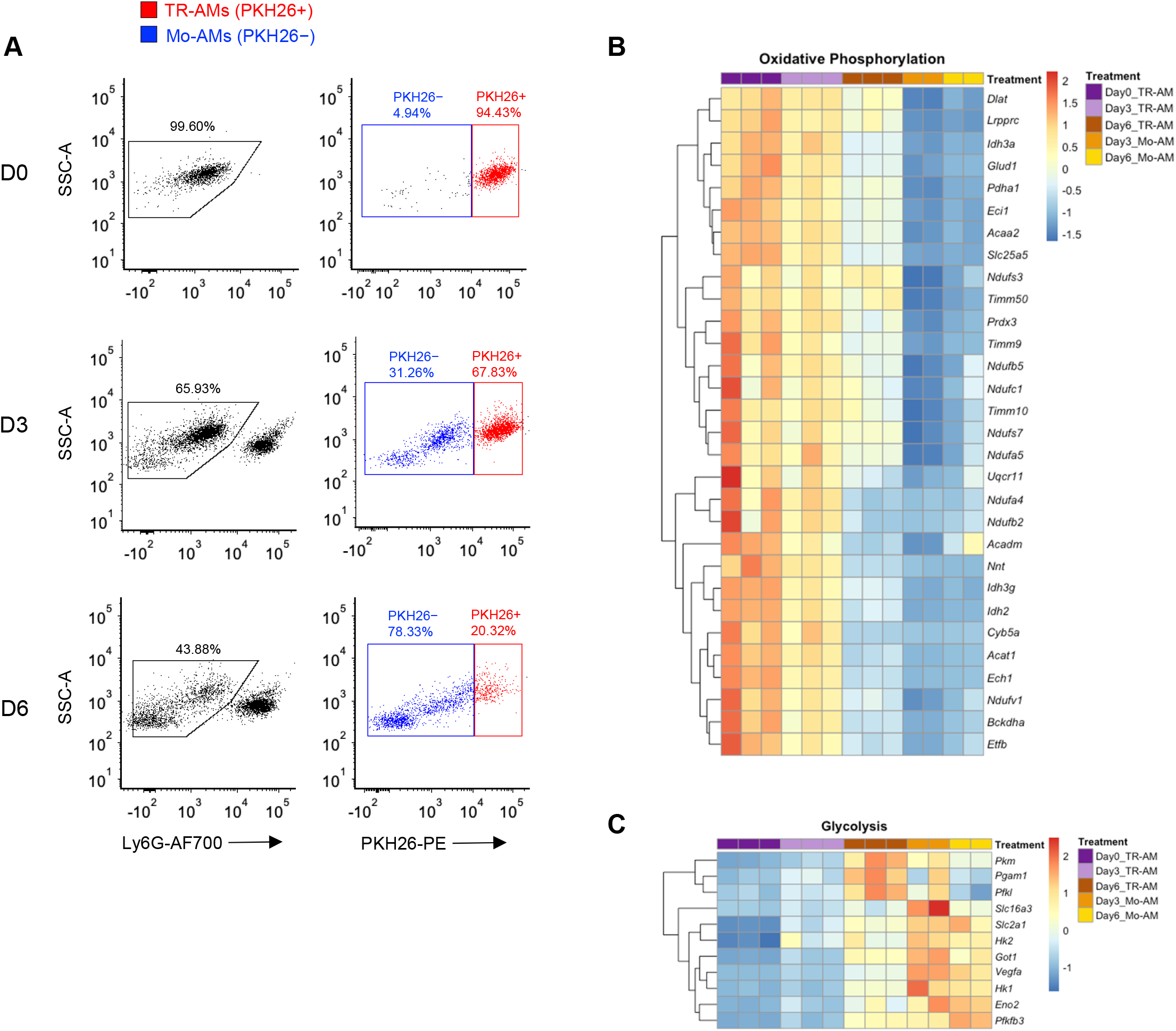
TR-AM survival correlates with a shift to glycolytic metabolism during influenza-induced acute lung injury. **(A)** FACS plots of BALF samples collected from C57BL/6 mice infected with PR8 (100 PFU) at baseline (D0), 3 days (D3) and 6 days (D6) post infection. First, debris, red blood cells, and lymphocytes were eliminated based on size (FSC) and granularity (SSC). Samples were first gated on single cells based on the side scatter (SSC)/forward scatter signal (FSC), and then live cells were selected (SYTOX Green-). Ly6G- used to exclude neutrophils. TR-AMs were identified as being PKH26+, and nonresident/infiltrating Mo-AMs were PKH26-. Gene expression heatmaps representing **(B)** oxidative phosphorylation and **(C)** glycolytic gene expression. Heatmaps were generated through DEG analysis of UniProt oxidative phosphorylation and glycolysis gene sets for FAC TR-AMs (PKH+; n=3/group) and Mo-AMs (PKH26-; n=2/group) over the infection time course.

### HIF-1α stabilization increases TR-AM survival and improves outcomes in influenza-induced acute lung injury

Given that the reduced TR-AM population on D6 presented with a glycolytic gene signature, we hypothesized that TR-AM survival was dependent upon a metabolic shift to glycolysis. In other words, a decrease in TR-AM numbers overtime was due to a large fraction of the cells failing to metabolically adapt to the conditions of the infected/hypoxic alveoli. Thus, TR-AMs that could not adapt to hypoxia and retained primarily mitochondria-driven metabolism died off while TR-AMs that shifted to glycolytic metabolism survived. To test this hypothesis, we first sought to determine if stabilization of HIF-1α was sufficient to induce a hypoxic metabolic state in AMs without altering O_2_ levels. To do this, we treated cells with FG-4592, an inhibitor of HIF prolyl hydroxylases. FG-4592 has a greater potency and fewer off target effects compared to DMOG, which broadly inhibits 2-oxoglutarate-dependent oxygenases (Singh, Wilson et al. 2020). TR-AMs treated with FG-4592 for 16 hours experienced a significant dose-dependent increase in glycolysis. (Figure 7A, B). FG-4592 induced robust HIF-1α stabilization leading to increased expression of HK2, LDHA, PHD1, and PHD3 (Figure 7C, D). Unlike hypoxia (1.5% O_2_), FG-4592 had very little impact on overall mitochondrial fitness in TR-AMs (Figure 7E). Basal respiration and mitochondrial ATP production were reduced, but spare mitochondrial compacity was increased, signaling a shift toward glycolytic ATP production at baseline but no loss in overall mitochondrial function (Figure 7E–G). Like hypoxia, FG-4592 treatment could also rescue ETC inhibitor-induced impairment in cytokine production (Figure 7H) and cell death in TR-AMs (Figure 7I). However, unlike TR-AMs exposed to 1.5% O_2_, FG-4592 did not broadly alter LPS cytokine responses suggesting that changes in the cytokine profile under hypoxia are oxygen-dependent, but remain independent of HIF-1α stabilization (Figure 7H).

**Figure 7.**
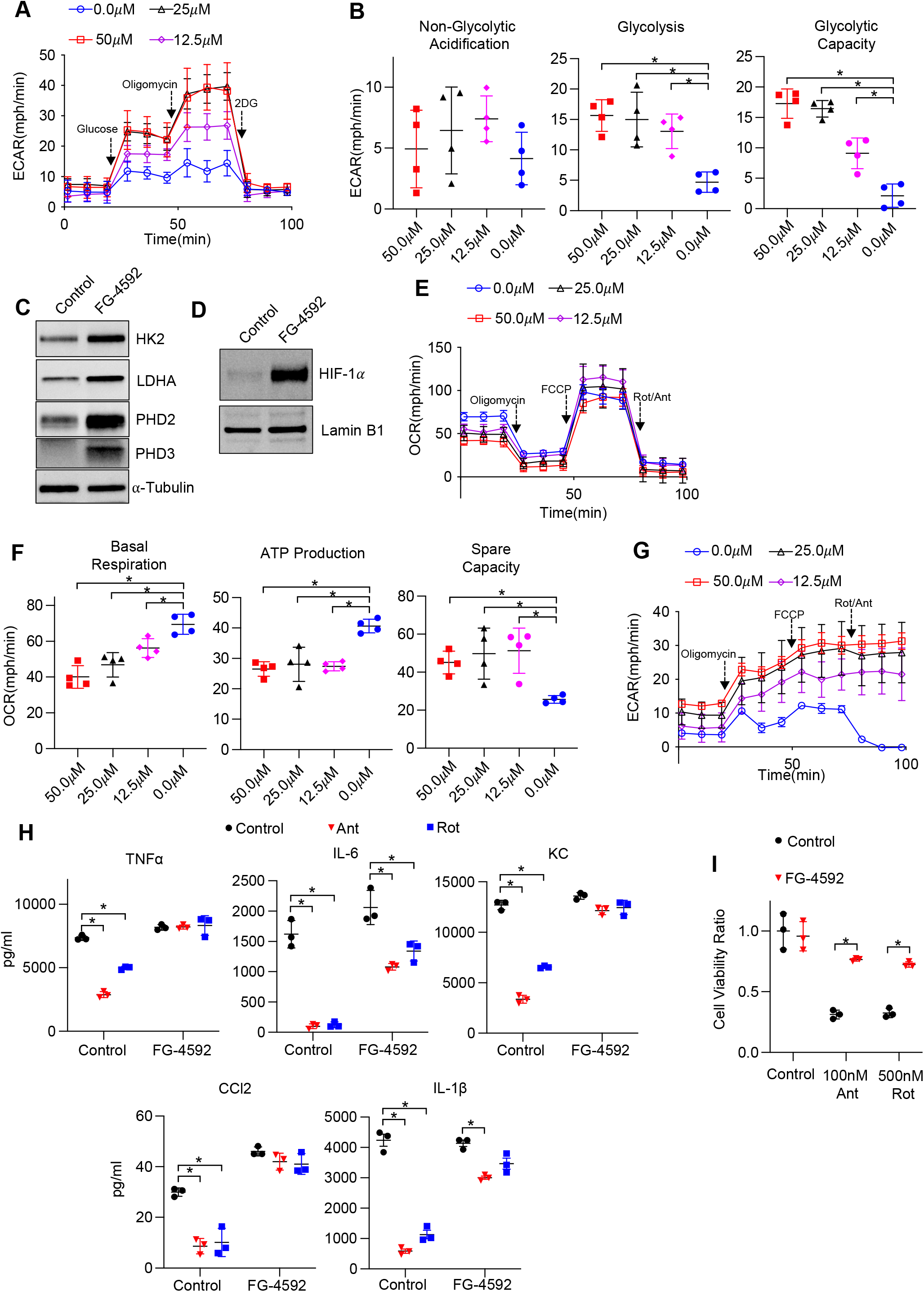
Non-hypoxic stabilization of HIF-1α increases TR-AM survival and improves outcomes in influenza-induced acute lung injury. TR-AMs were treated (16h) overnight ± FG-4592 (25.0*μ*M when not stated otherwise). **(A) G**lycolysis was measured as extracellular acidification rate (ECAR). **(B)** Quantification of glycolytic parameters. Data represent at least 3 independent experiments (n=4 separate wells per group). Glycolytic parameters compared to control group (0.0*μ*M) and significance was determined by one-way ANOVA with Bonferroni’s post test **(C)** Western blot analysis of whole cell lysate. **(D)** nuclear extract**. (E)** Mitochondrial stress test to measure oxygen consumption rate (OCR). **(F)** Quantification of mitochondrial respiration parameters. Data represents at least 3 experiments (n=4 separate wells per group). Mitochondrial parameters were compared to control group (0.0*μ*M) and significance was determined by one-way ANOVA with Bonferroni’s post test **(G)** ECAR measurement during mitochondrial stress test. **(H)** TR-AMs were pretreated overnight (16h) with 0.0*μ*M (no treatment) or 25.0*μ*M FG-4592 then stimulated with 20ng/ml LPS in the presence of absence of mitochondrial inhibitors (20nM Antimycin A (Ant) or Rotenone (Rot)) for 6 hours while maintaining pretreatment conditions. Sandwich ELISA was used to measure secreted cytokine (TNFα, IL-6, KC and CCL-2). Data represents at least 3 independent experiments; n=3 per group. **(I)** TR-AMs were treated with FG-4592 for 6h then treated with mitochondrial inhibitors (100nM Ant or 500nM Rot) overnight and an SRB assay was performed to measure cytotoxicity. Bar graphs represent cytotoxicity compared to control, 0.0uM group. Data represents at least 3 independent experiments (n=3 per group). Significance was determined by two-way ANOVA with Bonferroni’s post test. All error bars denote mean ± SD. *, p < 0.05

We next treated mice intratracheally with one dose of FG-4592 at the time of infection to evaluate the effect of early glycolytic adaptation on TR-AM survival and influenza-induced acute lung injury. Strikingly, FG-4592 treatment resulted in increased TR-AM (PKH26+ cells) survival at 6 days post infection (dpi) compared to infected controls (Figure 8A). The increase in TR-AM survival in FG-4592-treated mice was associated with reduced alveolar permeability (Figure 8B). FG-4592 treatment also led to a reduction in pro-inflammatory cytokine levels within the alveolar space at 6dpi (Figure 8C). Most importantly, FG-4592 treated mice experienced reduced weight loss and improved survival compared to infected controls (Figure 8D, E). Taken to together, these data suggest intratracheal FG-4592 treatment can increase TR-AM survival and improve outcomes in influenza-infected mice.

**Figure 8.**
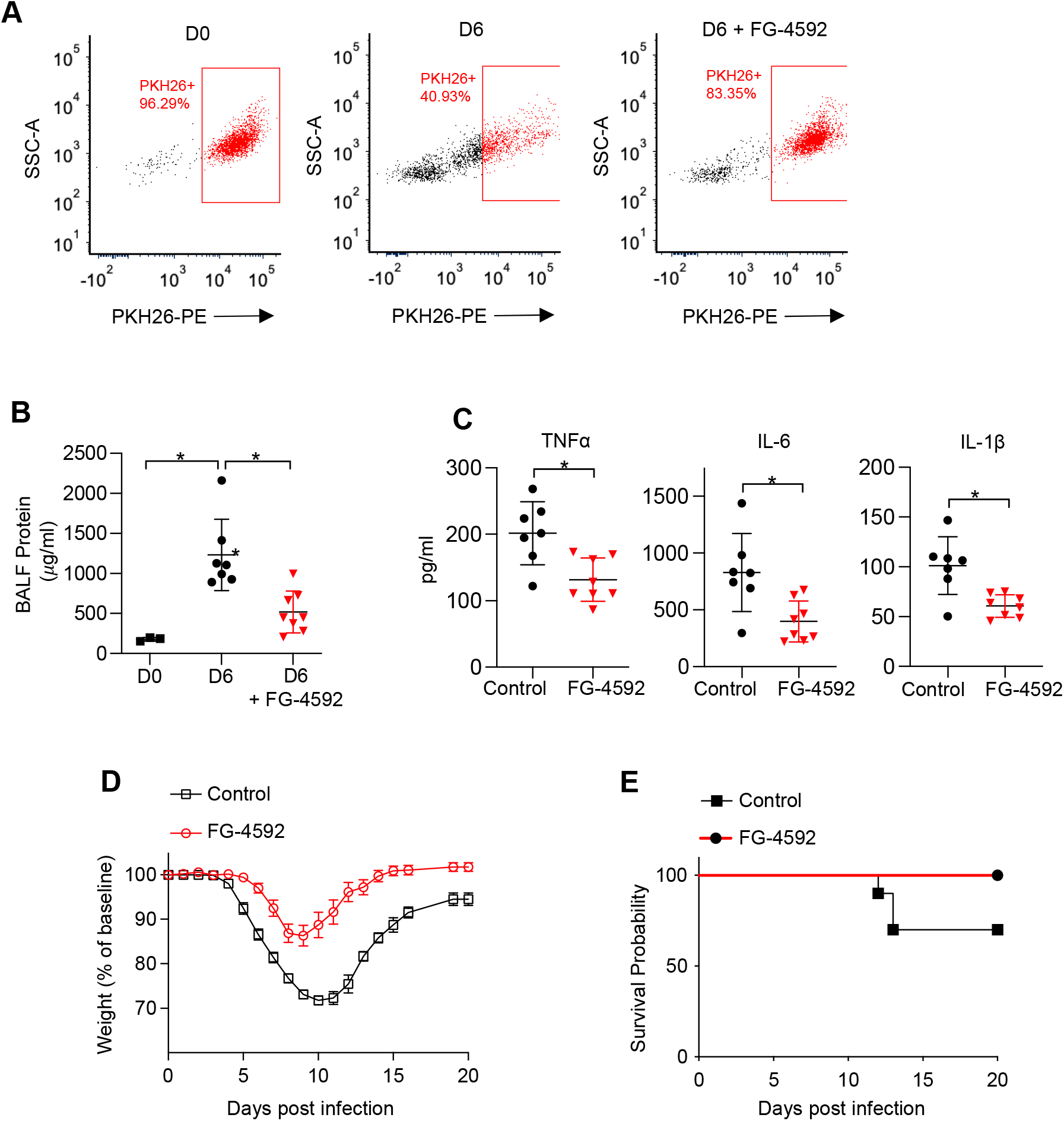
Non-hypoxic stabilization of HIF-1α increases TR-AM survival and improves outcomes in influenza-induced acute lung injury. We intratracheally infected C57BL/6 mice with PR8 (100 pfu) and collected bronchoalveolar lavage fluid (BALF) on day 0 (D0) (uninfected) and day 6 (D6) post-infection. Mice also received either the HIF-1α stabilizer (FG-4592) or vehicle control on D0. **(A)** Representative FACS plot of BALF macrophages. **(B)** BALF protein concentration **(C)** BALF proinflammatory cytokine levels at D6. BALF data generated from 2 separate experiments (n=7 mice/control group and n=8 mice/FG-4592 group) BALF data significance was determined by two-tailed Student’s t test. **(D,E)** C57BL/6 mice infected with PR8 (200pfu) (10 mice/group). **(D)** Weight loss represented as percentage and normalized to D0. **(E)** Survival curve. *p < 0.05. All error bars denote mean ± SD.

## DISCUSSION

ARDS is associated with high morbidity and mortality. Despite many decades of research, treatment remains supportive and there is no therapy that directly targets the pathogenesis of ARDS. Infection is the main cause of ARDS. Respiratory viruses such as influenza A virus and SARS-CoV-2 cause significant mortality by causing ARDS. In fact, the 2009 influenza pandemic and the ongoing COVID19 pandemic have shown that acute respiratory illnesses can have a profound effect on society in the 21^st^ Century. ARDS caused by influenza A and SARS-CoV-2 viruses is associated with loss of TR-AMs, that correlates with disease severity and mortality (Ghoneim, Thomas et al. 2013, Liao, Liu et al. 2020, Grant, Morales-Nebreda et al. 2021, Zhu, Wu et al. 2021). Because TR-AMs maintain a central role in lung homeostasis and response to airborne pathogens, understanding basic TR-AM processes, like metabolic adaptation in response to an altered lung environment during acute lung injury/ARDS, may allow us to therapeutically rescue TR-AM cell death, and augment their function to improve outcomes in acute respiratory illnesses.

We have previously demonstrated that TR-AMs are remarkably adapted to the high oxygen, low glucose environment of the alveolar lumen and do not require glucose or glycolysis to carry out their effector function as monocyte-derived macrophages do (Woods, Kimmig et al. 2020). ARDS leads to significant hypoxia to which TR-AMs need to adapt; thus, we sought to determine whether TR-AMs displayed metabolic plasticity during conditions of low oxygen. In this study, we found that TR-AMs adapted to hypoxia through HIF-1α induction. Hypoxia stabilized TR-AM HIF-1α in a dose-dependent manner resulting in robust increases in glycolytic protein expression and function. These changes were dependent on HIF-1α, as hypoxic TR-AMs treated with echinomycin, an inhibitor of HIF-1α DNA binding activity, lost their glycolytic capabilities. In contrast, BMDMs exhibited robust HIF-1α stabilization under normoxic conditions and BMDM HIF-1α expression and glycolytic output remained unchanged in response to hypoxia.

Transcriptomic analysis found 741 DEGs in TR-AMs in response to hypoxia compared to only 260 DEGs in BMDMs. Taken together, these data suggest that TR-AMs, although adapted to their unique high oxygen and low glucose environment, are capable of using glycolytic metabolism under stress conditions. Indeed, our *in vivo* experiments show that although TR-AMs are lost during ARDS, the surviving TR-AMs are glycolytically adapted and resemble recruited macrophages in metabolic gene expression. This is consistent with our previous findings that TR-AM viability is exquisitely sensitive to mitochondrial inhibition while BMDM viability is unaffected. Our current findings show that hypoxia restores viability to TR-AMs under mitochondrial inhibition and reveres the suppressive effects of this inhibition on TR-AM cytokine production supporting optimal effector function in a low oxygen environment. These data suggest that the link between immune effector function and metabolism in TR-AMs is based on overall cellular fitness, and that the environmental shift from high oxygen and low glucose under steady-conditions to low oxygen and increased glucose during lung injury may necessitate cellular adaptation to the microenvironment to ensure survival.

Pharmacological depletion of TR-AMs has offered insight into their beneficial immunoregulatory properties in various lung injury models. TR-AMs have been shown to alleviate lung injury by clearing apoptotic neutrophils, suppressing T-cell-mediated inflammatory responses, and limiting dendritic cell infiltration and antigen presentation (Thepen, Van Rooijen et al. 1989, Holt, Oliver et al. 1993, Knapp, Leemans et al. 2003, Jakubzick, Tacke et al. 2006). Loss of TR-AMs during influenza infection is known to enhance mortality, and several studies have shown that there is a progressive loss of TR-AMs over the time course of infection (Kim, Lee et al. 2008, Ghoneim, Thomas et al. 2013, Schneider, Nobs et al. 2014, Cardani, Boulton et al. 2017, Zhu, Wu et al. 2021). Why TR-AMs are lost during the course of IAV infection is not understood. We hypothesized that failure to adapt to the hypoxic environment may play a role in the TR-AM loss. We found that HIF-1α activation was sufficient to promote glycolysis, and rescue TR-AM viability and effector function under mitochondrial inhibition. Furthermore, when mice were treated with a HIF-1α stabilizer at the time of influenza infection, TR-AM survival was increased, and accompanied by reduced lung injury and death. These findings suggest that HIF-1α is essential for TR-AM cell survival, and that increasing TR-AM cell number by promoting their adaptation to ARDS associated changes in the microenvironment during infection reduces lung injury.

In agreement with our findings, Zhu and colleagues recently showed that TR-AMs at D6 of influenza infection have high HIF-1α expression; however, they suggested that HIF-1α stabilization in TR-AMs worsens lung injury through enhanced proinflammatory effector function (Zhu, Wu et al. 2021). This study relied on the use of a non-inducible *Cd11c* cre allele to knockout HIF-1α in TR-AMs. While TR-AMs do express high levels of Cd11c, this marker is not specific for TR-AMs, and is also expressed in different monocyte/macrophage populations, dendritic cells, and natural killer cells (Abram, Roberge et al. 2014). Moreover, as monocyte-derived macrophages enter the alveolar space, they begin to express Cd11c (Misharin, Morales-Nebreda et al. 2013). Thus, it cannot be determined from these experiments whether the observed effect of HIF-1α deletion on inflammation and lung injury was specific for TR-AMs as the recruited monocyte-derived macrophages will also lose HIF-1α as they enter the lungs. It is likely that the recruitment of monocyte-derived macrophages was inhibited by HIF-1α deletion, resulting in reduced inflammation and lung injury (Cramer, Yamanishi et al. 2003). Using FG-4592 *in vitro*, we found that HIF-1α stabilization and glycolytic reprogramming resulted in no significant changes in proinflammatory cytokine production downstream of LPS. Furthermore, we show that treatment of mice with FG-4592 promoted survival of TR-AMs after influenza infection, and lead to reduced levels of lung injury. An inducible model of HIF-1α activation will be required to conclusively demonstrate whether HIF-1α expression specifically in TR-AMs promotes survival during lung injury since constitutive activation of HIF-1α leads to defects in TR-AM development (Izquierdo, Brandi et al. 2018). There have also been no studies on the effects of long-term HIF-1α activation and resultant metabolic reprogramming in TR-AMs under steady state or during recovery from inflammatory stimuli. Thus, our short-term pharmacologic activation of HIF-1α and glycolytic reprogramming in TR-AMs is a useful model for studying the role of metabolic reprogramming during lung injury.

Interestingly, we found that the effects of hypoxia on both TR-AM and BMDM effector function appears to be independent of HIF-1α. While HIF-1α stabilization was sufficient to drive metabolic reprogramming in TR-AMs, FG-4592 did not affect cytokine production downstream of LPS. By contrast, when cultured in hypoxia TR-AMs exhibited marked changes in their secreted cytokine profile including increased secretion of TNF-α, KC, and IL-1β and less IL-6 and CCL2 compared to normoxic controls. BMDMs exposed to hypoxia and treated with LPS responded similarly to TR-AMs with the exception to CCL2, which remained unchanged in response to hypoxia in BMDMs. This was independent of any effect of hypoxia on HIF-1α stabilization or target gene expression in BMDMs. Similarities between TR-AMs and BMDMs in terms of cytokine profiles would suggest that low oxygen concentrations may be driving these changes independently of HIF-1α stabilization. Further investigation will be required to elucidate the mechanisms by which a low oxygen environment alters cytokine production.

In conclusion, under normoxic conditions, TR-AMs depend on mitochondrial respiration, inhibition of which leads to decreased effector response and cell death. Under hypoxic conditions, as occurs during ARDS, TR-AMs stabilize HIF-1α in a dose dependent manner and have a robust HIF-1α response compared to BMDMs. Stabilization of HIF-1α allows TR-AMs to augment glycolytic function, and prevents their death following the inhibition of mitochondrial respiration as well as following influenza A viral infection. These data suggest that therapies inducing HIF-1α in TR-AMs may be beneficial in ARDS by preventing their death through metabolic adaptation to the ARDS microenvironment that is low in O_2_ and high in glucose.

## MATERIALS AND METHODS

### Primary Culture of Macrophages

All studies in animals were approved by the Institutional Animal Care and Use Committee at the University of Chicago IACUC. 6-8 week old C57BL/6 mice were humanely euthanized, and their TR-AMs were isolated via standard bronchoalveolar lavage (intratracheal instillation) using PBS + 0.5 mM EDTA. Following isolation, TR-AMs were counted, plated in RPMI 1640 (ThermoFisher, catalog number 11875119) supplemented with 10% FBS (Gemini, catalog number 100-106) and 1% penicillin-streptomycin (Gemini, catalog number 400-109), and allowed to adhere to tissue culture plates for one hour prior to experimentation. BMDMs were generated by isolating bone marrow cells from the femur and tibia bones of 6-8 week old C57BL/6 mice. Bone marrow cells were differentiated into BMDMs using 40 ng/mL recombinant M-CSF (BioLegend, catalog number 576406) in the same media formulation as TR-AMs. On day seven, BMDMs were replated and allowed to adhere to tissue culture plates for two hours prior to experimentation. After adherence, cells were washed, fresh media added, and placed under experimental conditions. For inflammatory stimulation, LPS was used at a concentration of 20ng/ml. For hypoxia experiments, macrophage cultures were placed in an airtight incubator system that utilizes N_2_ displacement of O_2_ to achieve hypoxic conditions (Coy Hypoxic Chamber-O_2_ Control InVitro Glove Box). The sealed system ensures minimal fluctuations in O_2_ levels in experiments when treating cultures and collecting samples under hypoxic conditions.

### Bioenergetic Measurements

Glycolytic and mitochondrial respiration rates were measured using the XFe24 Extracellular Flux Analyzer (Agilent, Santa Clara, MA). BMDMs and TR-AMs were seeded at 4.0 × 10^4^/well onto Seahorse XF24 Cell Culture Microplates. Cells were equilibrated with XF Base media (Agilent, catalog number 103334-100) at 37 °C for 30 minutes in the absence of C02. Glycolytic rate was assessed using the manufacturers’ protocol for the Seahorse XF Glycolysis Stress Test followed by sequential injections with glucose (10mM), oligomycin (1.0μM), and 2-DG (100mM).

Mitochondrial respiration rate was measured using the Seahorse XF Mito Stress Test according to the manufacturer’s protocol followed by sequential injections with oligomycin (1.0μM), FCCP (1.0μM for BMDMs and 4.0μM for TR-AMs), and rotenone/antimycin A (1.0μM). Assessment of real-time metabolic responses to LPS was performed using the protocol detailed in an application note provided by the Agilent (Kam Y 2017). In brief, following plating, cells were equilibrated in XF base media supplemented with 10 mM glucose, 2 mM L-glutamine, 1 mM sodium pyruvate (Sigma, catalog number 11360070) and 5 mM HEPES (Sigma, catalog number 15630080), pH 7.4 and incubated at 37 °C without CO2 for 30 minutes prior to XF assay. Baseline metabolic rates were measured followed by direct injection of LPS (final concentration:20ng/ml). Bioenergetic rates were subsequently measured every three minutes for approximately 5 hours in total.

Due to the limitations of the XFe24 Extracellular Flux Analyzer, all bioenergetic analysis on hypoxic samples was performed in the following manner. Cells were treated under hypoxic conditions (most commonly for 16 hours) then bioenergetic analysis was performed under normoxic conditions. Moreover, it is impossible to evaluate varying levels of hypoxia on a single Seahorse microplate. Thus, energy curves comparing varying levels of O_2_ (i.e. Figure 1A) were performed individually and then subsequently represented on the same graph for comparison. All individual experiments were repeated a minimum of three times to an ensure accurate representation and statistical comparison.

### Cell lysis, subcellular fractionalization and Immunoblotting

Whole cell lysates were prepared by scraping cells into lysis buffer containing 25mM Tris·HCl (pH 7.6), 150mM NaCl, 1% NP-40, 1% sodium deoxycholate, 0.1% SDS, 0.1% Benzonase, and Halt™ Protease Inhibitor Cocktail (ThermoFisher, catalog number 1861284 and 78430). Samples were centrifuged at 16,000 x *g* at 4 °C for 5 min to pellet cellular debris. Subcellular fractionalization and lysate preparation were carried out using the NE-PER Nuclear and Cytoplasmic Extraction Reagents (ThermoFisher, catalog number 78833). Lysate protein concentration was determined using the Pierce™ BCA Protein Assay Kit (ThermoFisher, catalog number 23225). Equal concentrations of samples (15μg for whole cell lysates and 5μg for nuclear fractions) were resolved on Criterion gels (Bio-Rad, catalog number 5671093 and 5671094) and transferred to nitrocellulose (Bio-Rad, catalog number 1620167). Primary antibodies used were rabbit anti-HK2 (Cell signaling, catalog number C64G5, 1:1000), rabbit anti-LDHA (Cell signaling, catalog number 20125, 1:1000), rabbit anti-PHD2/Egln1 (Cell Signaling, catalog number 4835, 1:1000), rabbit anti-Egln3/PHD3 (Novus Biologicals, catalog number NB100-303, 1:1000), mouse anti-IL1β (Cell signaling, catalog number 12242 1:1000), rabbit anti-Lamin B1 (Proteintech, catalog number 12987-1-AP, 1:1000), rabbit anti-HIF-1α (Caymen Chemical, catalog number 10006421, 1:500), and mouse anti-tubulin (Sigma, catalog number T6074, 1:20,000). Secondary antibodies used were anti-rabbit IgG HRP-linked antibody (Cell Signaling, catalog number 7074, 1:2,500) and goat anti-mouse IgG HRP-linked antibody (Cell Signaling, catalog number 7076, 1:2,500). Protein expression was visualized using Immobilon ECL Ultra Western HRP Substrate (Millipore Sigma, catalog number WBULS0500) in combination with the BioRad ChemiDoc Touch Imaging system. All immunoblot data were repeated in at least three independent experiments.

### Quantitative PCR

RNA was isolated from cells using the Direct-zol RNA MiniPrep kit (Zymo Research, catalog number R2052) and reverse transcribed using iScript Reverse Transcription Supermix (Bio-Rad, catalog number 1708841). Quantitative mRNA expression was determined by real-time RT-PCR using iTaq Universal SYBR Green Supermix (Bio-Rad, catalog number 172-5121). *rlp19* served as a housekeeping gene, and gene expression was quantified using the ΔΔct method to determine relative fold-change The following mouse-specific primer sequences were used: *rlp19* (5’-CCGACGAAAGGGTATGCTCA-3’, 5’-GACCTTCTTTTTCCCGCAGC-3’), *Il6* (5’-TTCCATCCAGTTGCCTTCTTGG-3’, 5’-TTCCTATTTCCACGATTTCCCAG-3’), *tnfα* (5’-AGGGGATTATGGCTCAGGGT-3’, 5’-CCACAGTCCAGGTCACTGTC-3’, *il1* (5’-GCCACCTTTTGACAGTGATGAG, 5’-GACAGCCCAGGTCAAAGGTT-3’), *kc* (5′-AGACCATGGCTGGGATTCAC-3′, 5′-ATGGTGGCTATGACTTCGGT-3′), *ccl2* (5’-CTGTAGTTTTTGTCACCAAGCTCA-3’, 5’-GTGCTGAAGACCTTAGCCCA-3’).

### RNA-Sequencing

RNA was isolated and submitted to the University of Chicago Genomics Core Facility for sequencing with the Illumina NovaSEQ6000 sequencer (100bp paired-end). Sequencing read (FASTQ) files were generated and assessed for per base sequence quality using FastQC. Reads were mapped to the mouse genome (GRCm38.p6, GENCODE) using Spliced Transcripts Alignment to a Reference (STAR) software, and the resulting gene transcripts were quantified using featureCounts.

Gene counts were then imported into R for differential expression analysis using the Bioconductor package DESeq2. Gene counts were filtered to remove low-expressing genes at a threshold of 2 counts per million. Differential expression was calculated between normoxia and hypoxia groups for both AMs and BMDMs. Differential gene expression was considered significant for genes with an FDR-adjusted p-value < 0.05 and fold change (FC) > 2. Reactome enrichment hit pathways and the linked gene lists from significant DEGs were identified by using Reactome Cytoscape Plugin (Shannon, Markiel et al. 2003, Wu, Feng et al. 2010). Oxidative phosphorylation and glycolysis gene sets were extracted from UniProtKB, then their associated DEG read counts (TPM) were normalized using gensvm R package gensvm.maxabs.scale function and center scaled for heat map visualization. All heatmaps were generated with Pretty heatmaps R package pheatmap function.

### Cytokine Analysis

Secreted TNFα, IL-6, KC, CCl2, and IL-1β levels were evaluated in macrophage media using a standard sandwich ELISA (R&D Systems DuoSet ELISA Development System, catalog numbers DY410, DY406, DY453, DY479 and DY401). For IL-1*β* sample collection, 5mM ATP was added to TR-AM cultures for 30 minutes following 6h LPS treatment to activate caspase 1, ensuring proIL-1*β* cleavage and IL-1β release. Rotenone and Antimycin A concentrations were 20nM for TR-AMs and 1μM for BMDMs when used in ELISA experiments.

### Metabolomics

TR-AMs were plated at 2.5 × 10^6^ on 60 mm plates for metabolite extraction. Following treatment, cells were washed twice with a 5% mannitol solution and metabolites were extracted using 400μl 100% methanol. 275μl of aqueous internal standard solution was mixed in with the methanol and the extract solution was transferred to a microcentrifuge. The extracts underwent centrifugation at 2,300 x *g* at 4 °C for 5 min to precipitate insoluble material and the resulting supernatant was transferred to centrifugal filter units (Human Metabolome Technologies (HMT), Boston, MA). Filtering of supernatant occurred at 9,100 x *g* at 4 °C for 2 hours. The filtrate was sent to HMT and analyzed using capillary electrophoresis–mass spectrometry.

### Sulforhodamine B (SRB) Colorimetric Assay

In vitro cytotoxicity was measured using the SRB assay (Vichai and Kirtikara 2006). Following treatment, cells were fixed in 10% TCA and then stained with SRB dye. Cellular protein-dye complexes were solubilized in 10mM Tris base and the samples were read at OD 510 using a microplate reader. Data was normalized to the untreated, normoxia groups, which were representative of no cellular damage. ETC inhibitor concentration in BMDMs were as follows: 1μM rotenone, and 1μM antimycin A. ETC inhibitor concentration in TR-AMs were as follows: 500nM rotenone, and 100nM antimycin A.

### Lactate Assay

Secreted lactate was measured using the lactate colorimetric assay kit (Sigma, catalog number MAK064-1KT). Cells were plated in serum-free DMEM media (RPMI media and serum interfere with assay), allowed to adhere, washed with phosphate buffer saline, and exposed to normoxia or 1.5% O_2_. Samples were collected at 16 hours post treatment and manufacturer’s protocol was followed to measure lactate.

### Murine Influenza infection protocol and Survival Studies

C57BL/6 mice (6–8 weeks old) were anesthetized and challenged intratracheally (IT) with mouse-adapted influenza (A/PR8/34; 200 plaque-forming units [pfu]). A single FG-4592 (50uM) treatment was administered (IT) simultaneously with IAV. Body weight and survival was monitored every 24 hours for 20 days (10 mice/group). Body weight is represented as percent deviation from baseline at time of infection. “Influenza A virus, A/PR8/34 (H1N1), NR-348” was obtained through BEI Resources, NIAID, NIH.

### BALF analysis

C57BL/6 mice were euthanized and a single 0.5ml saline wash was instilled into the lungs via the trachea and subsequently collected. BALF protein concentration was determined using the Pierce™ BCA Protein Assay Kit (ThermoFisher, catalog number 23225). BALF TNFα, IL-6, and IL-1β were measured using sandwich ELISA.

### Flow Cytometry

C57BL/6 mice (6–8 weeks old) were anesthetized with isoflurane and underwent retro-orbital injection with 100 μl PKH26 Red Fluorescent Cell Linker Dye for Phagocytic Cell Labeling (catalog number PKH26PCL-1KT; Millipore Sigma) 1 day before lung challenge. The mice were then challenged intratracheally with IAV 100 (pfu). FG-4592 (50μM) was administered intratracheally at the same time of PR8 infection. After challenge, the mice were euthanized and immune cells were collected via BAL. BAL cells were first treated with Fc Block (clone 2.4G2, catalog number 553141; BD Biosciences) and stained with fluorochrome-conjugated antibodies. The antibodies used were AlexaFluor 700 anti-mouse Ly-6G (Clone 1A8, catalog number 127621, 1:250; BioLegend). Immediately before sorting, cells were resuspended in sorting buffer (0.2% BSA in PBS) containing 5 nM SYTOX Green Nucleic Acid Stain (catalog number S7020; ThermoFisher) to distinguish between live and dead cells. Cell sorting was performed on a FACS Aria II instrument and data were acquired using BDFACS Diva software and analyzed with FCS Express 7 software. First, debris, red blood cells, and lymphocytes were eliminated based on size (FSC) and granularity (SSC). Next gates selected for live cells (FITC−) and eliminated neutrophils (Ly6G+). Based on previous validation experiments, the remaining cells are of macrophage lineage with TR-AMs being PKH26_+_ (SiglecF_+_, F4/80_+_, Cd11c^Hi^, Ly6c^Lo^) and Mo-AMs being PKH26_+_ (F4/80, Cd11c^Lo^, Ly6c^Hi^). PKH26_+_ and PKH26_−_ cells were sorted into RNA lysis buffer and samples were prepared for RNAseq.

### Statistics

The data were analyzed in Prism 8 (GraphPad Software Inc.). All data are shown as mean ± SD. ANOVA was used for statistical analyses of data sets containing more than two groups, and Bonferroni’s *post hoc* test was used to explore individual differences. Statistical significance was defined as *P* <0.05.

---

**Figure S1.**
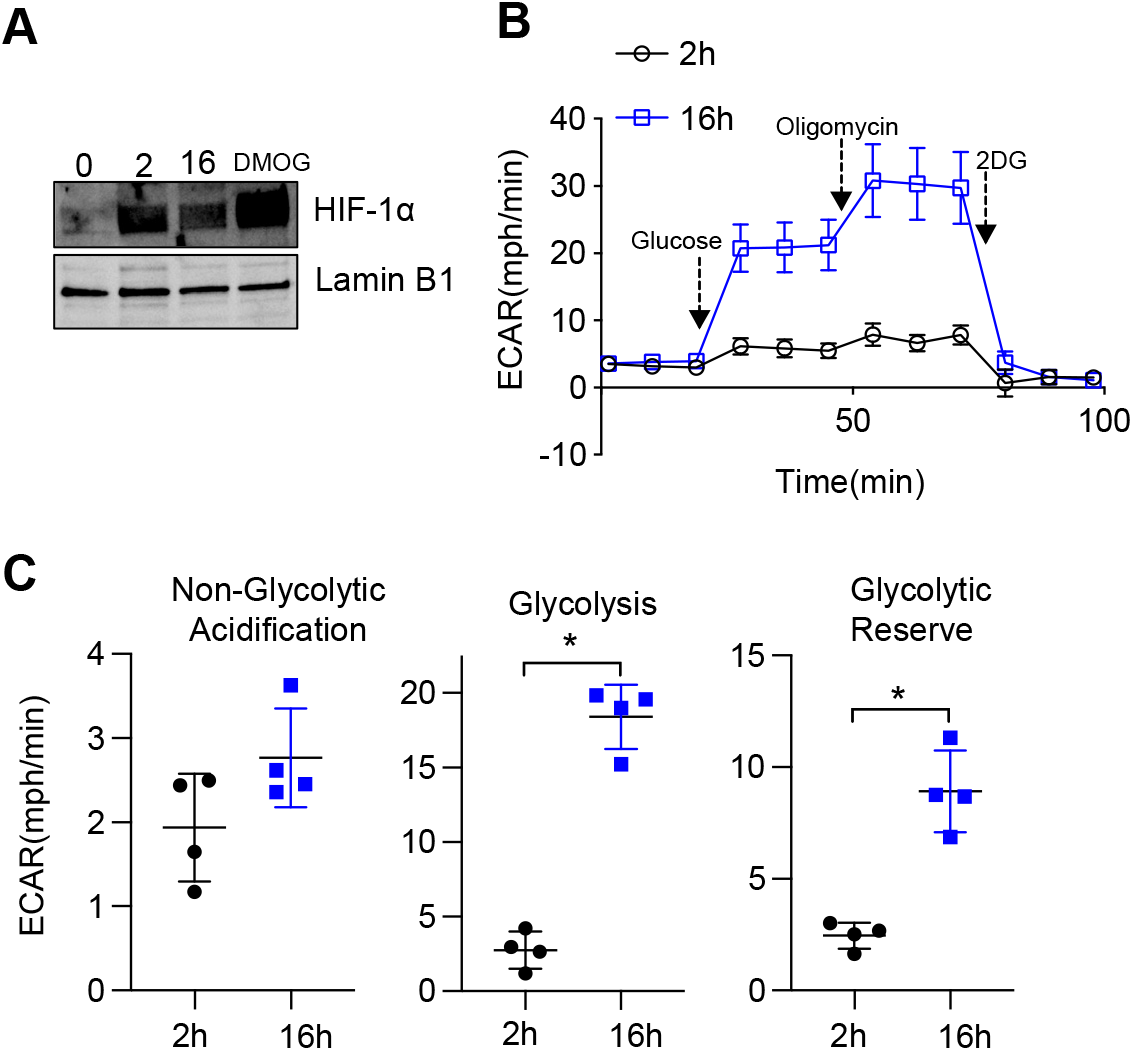
Prolonged but not short-term hypoxia induces glycolysis in TR-AMs. TR-AMs were incubated for 2h or overnight (16h) at 1.5% O_2_. **(A)** Western blot analysis of nuclear extracts to assess HIF-1α protein expression. **(B)** Using Seahorse XF24 technology, glycolysis was measured as extracellular acidification rate (ECAR). **(C)** Interleaved scatter plots quantifying glycolytic parameters. Data represents at least 3 independent experiments (n=4 separate wells per group). Significance was determined by two-tailed Student’s t test. *,p< 0.05.

**Figure S2.**
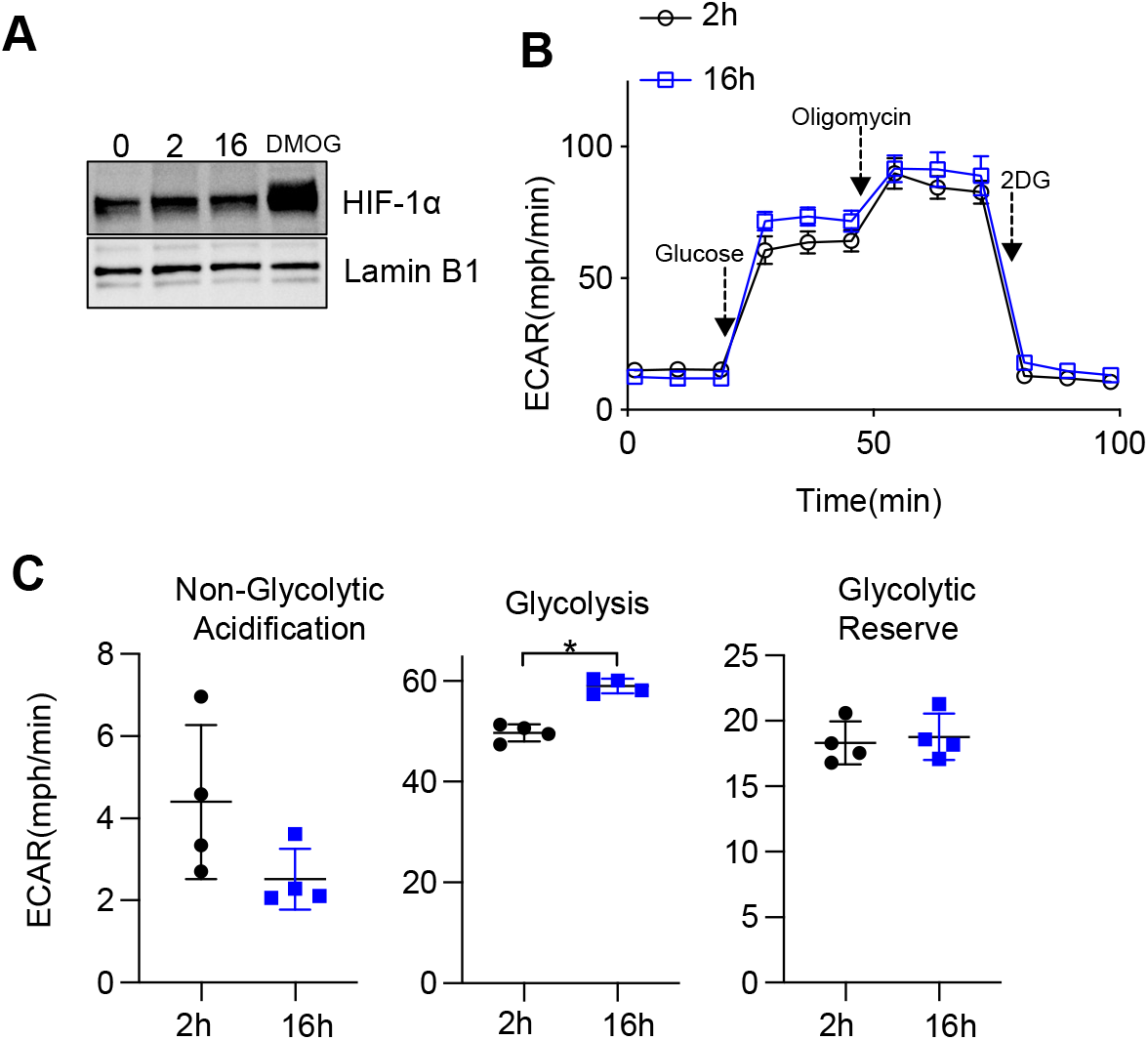
Short-term or prolonged hypoxia has no significant effect on HIF-1α expression or glycolysis in BMDMs. BMDMs were incubated for 2h or overnight (16h) at 1.5% O_2_. **(A)** Western blot analysis of nuclear extracts to assess HIF-1α protein expression. **(B)** Using Seahorse XF24 technology, glycolysis was measured as extracellular acidification rate (ECAR). **(C)** Interleaved scatter plots quantifying glycolytic parameters. Data represents at least 3 independent experiments (n=4 separate wells per group). Significance was determined by two-tailed Student’s t test. *, p < 0.05

**Figure S3.**
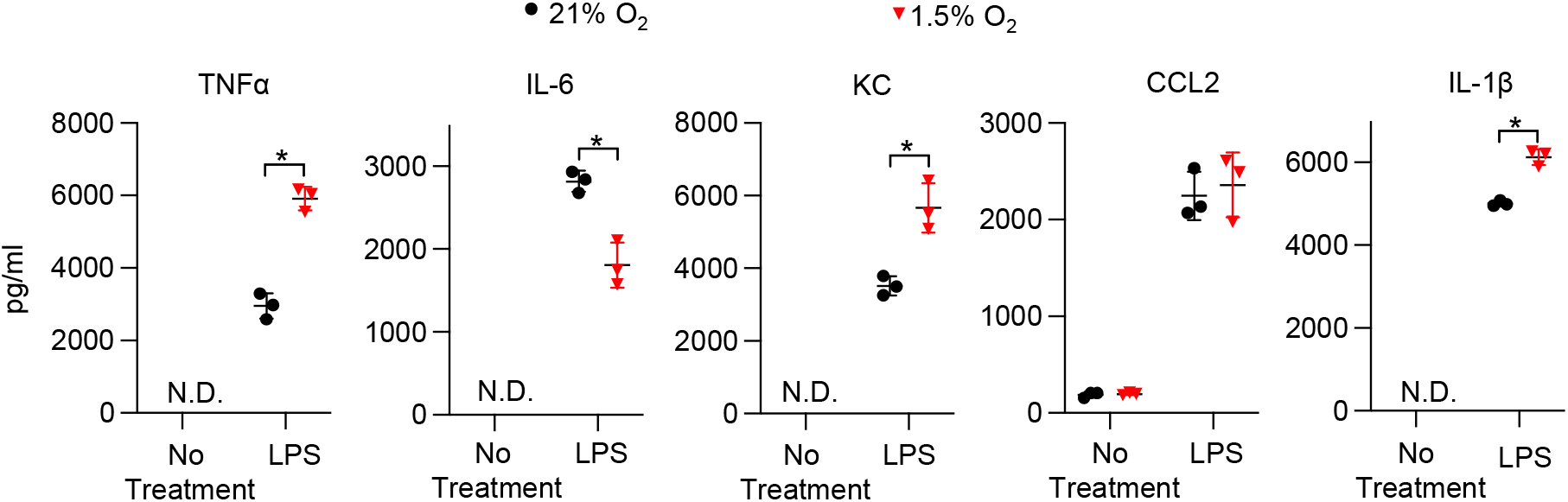
Hypoxia alters cytokine production in BMDMs. BMDMs were incubated overnight (16h) under normoxia or 1.5% O_2_ then stimulated with 20ng/ml LPS for 6 hours while maintaining pretreatment conditions. For IL-1*β*, 5mM ATP was added to BMDMs for 30 minutes following 6h LPS treatment to activate caspase 1, ensuring IL-1*β* release. **(A)** Sandwich ELISA was used to measure secreted cytokine (TNFα, IL-6, KC, CCL2 and IL-1*β*). Data represents at least 3 independent experiments; n=3 per group. Significance was determined by two-tailed T Test. All error bars denote mean ± SD. *, p < 0.05

**Figure S4.**
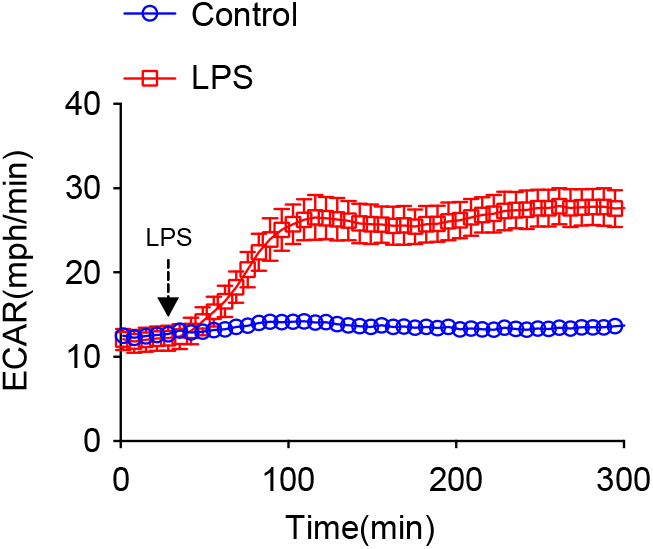
LPS induces an immediate increase in glycolysis in BMDMs. ECAR was measured in normoxic BMDMs following acute LPS injection (final concentration: 20 ng/ml).

**Figure S5.**
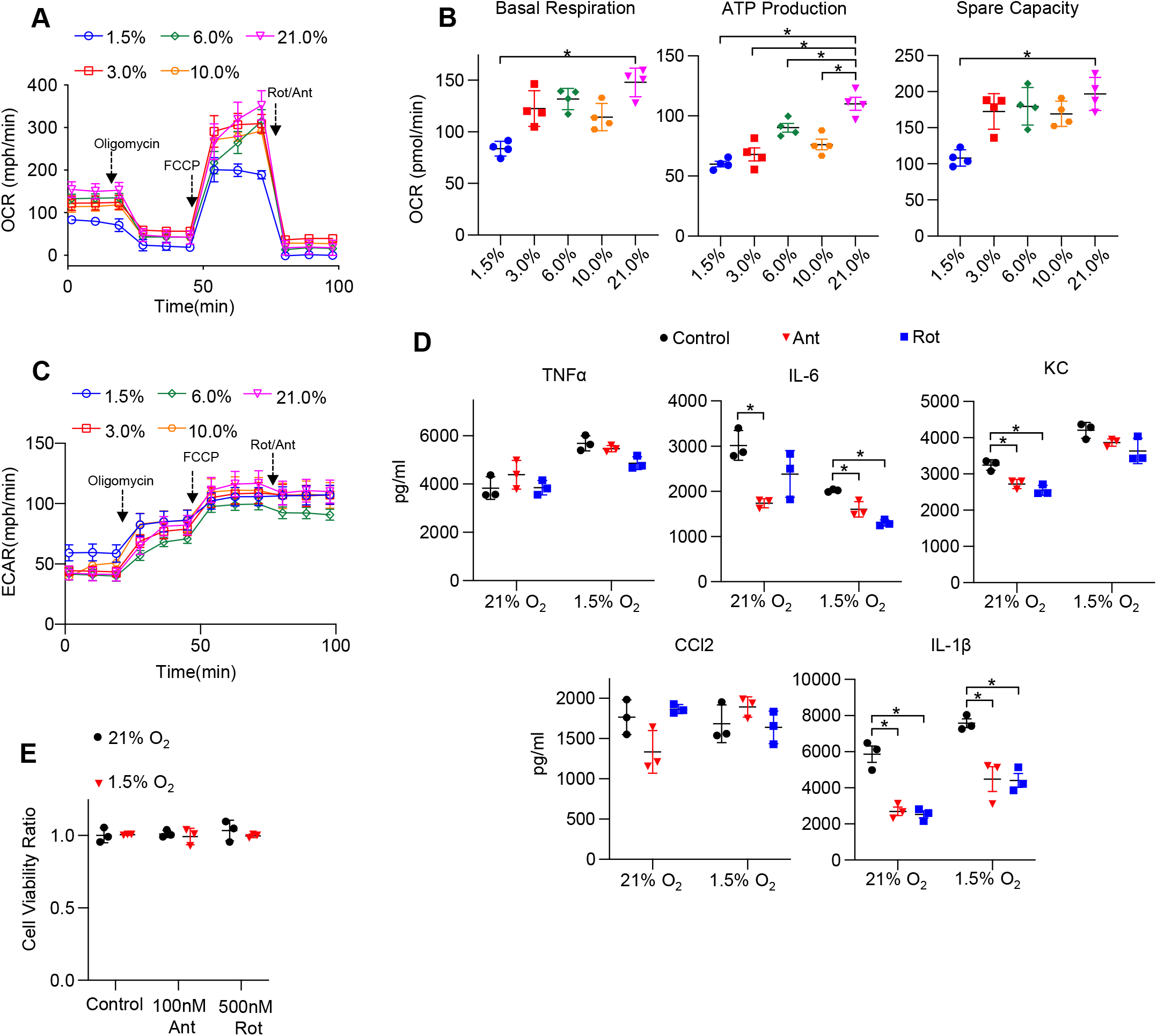
The effect of hypoxia on BMDM mitochondrial function, cytokine production and cell liability under ETC inhibition. **(A)** Mitochondrial stress test to measure oxygen consumption rate (OCR) using Seahorse XF24 in BMDMs. **(B)** Interleaved scatter plots quantifying mitochondrial respiration parameters. Data represents at least 3 experiments (n=4 separate wells per group). Mitochondrial parameters were were compared against 21% O_2_ and significance was determined by one-way ANOVA with Bonferroni’s post test **(C)** ECAR measurement during mitochondrial stress test. **(D)** BMDMs were incubated overnight (16h) under 21% or 1.5% O_2_ then stimulated with 20ng/ml LPS in the presence of absence of mitochondrial inhibitors (20nM Antimycin A (Ant) or Rotenone (Rot)) for 6 hours while maintaining pretreatment conditions. ELISA was used to measure secreted cytokine (TNFα, IL-6, KC, CCL2, and IL-1β) levels in media. ATP added to cells prior to collection for IL-1β assessment. Data represent at least 3 independent experiments; n=3 per group. Significance was determined by one-way ANOVA with Bonferroni’s post test. **(E)** BMDMs were cultured under 21% or 1.5% O_2_ for 6h then treated with mitochondrial inhibitors (100nM Ant or 500nM Rot) overnight and an Sulforhodamine B assay was performed to measure cytotoxicity. Graphs represent cell viability compared to control, 21% O_2_ group. Data represent at least 3 independent experiments (n=3 per group). Significance was determined by two-way ANOVA with Bonferroni’s post test. All error bars denote mean ± SD. *, p < 0.05.

